# Sensitive bacterial V_m_ sensors revealed the excitability of bacterial V_m_ and its role in antibiotic tolerance

**DOI:** 10.1101/2022.06.02.494477

**Authors:** Xin Jin, Xiaowei Zhang, Xuejing Ding, Tian Tian, Chao-Kai Tseng, Xinwei Luo, Xiao Chen, Chien-Jung Lo, Mark C. Leake, Fan Bai

**Author notes:** Correspondence to: Fan Bai. These authors contributed equally to this work (Xin Jin, Xiaowei Zhang, Xuejing Ding, Tian Tian).

## Abstract

As an important free energy source, the membrane voltage (V_m_) regulates many essential physiological processes in bacteria. However, in comparison with eukaryotic cells, knowledge of bacterial electrophysiology is very limited. Here, we developed a set of novel genetically encoded bacterial V_m_ sensors which allow single-cell recording of bacterial V_m_ dynamics in live cells with high temporal resolution. Using these new sensors, we reveal the electrically “excitable” and “resting” states of bacterial cells dependent on their metabolic status. In the electrically excitable state, frequent hyperpolarization spikes in bacterial V_m_ are observed, which facilitates increased antibiotic tolerance. In the electrically resting state, bacterial V_m_ displays significant cell-to-cell heterogeneity and is linked to the cell fate after antibiotic treatment.

## Introduction

Membrane voltage (V_m_) is an important free energy source for bacteria. It powers many fundamental physiological processes, such as oxidative phosphorylation (*1*), transmembrane transport (*2, 3*), flagellar motion (*4, 5*), and cell division (*6*). Since the bacterial V_m_ is tightly coupled to metabolism (*7*), V_m_ may also play critical roles in bacterial virulence and drug tolerance (*8–10*). It was generally believed that bacterial V_m_ is static, until recent studies revealed that bacterial V_m_ can be highly dynamic and enables cell-to-cell communication in biofilms (*7, 11, 12*). Kralj et al. reported the electrical spiking of transient V_m_ depolarization in *Escherichia coli* (*E. coli*) (*11*). It is also reported that the mechanosensation of *E. coli* is controlled by voltage-induced calcium flux (*13*). In *Bacillus subtilis* biofilm communities, long-range electrical signaling mediated by spatially propagating waves of potassium was found to coordinate metabolic states (*12*). Further research has shown that the electrical signaling generated by the biofilm can also attract distant motile cells, leading to incorporation of different species into a pre-existing biofilm community (*14*).

These recent findings necessitate new studies of bacterial electrophysiology (*7, 15*). However, the tools for monitoring bacterial V_m_ dynamics *in vivo* at the single-cell level are very limited (*7*). Due to the small size of bacteria and the existence of cell wall, the application of patch clamp, a gold standard for single-cell V_m_ tracking in eukaryotic cells, need to generate giant spheroplasts, which cannot reflect the normal physiology of bacteria (*15*). Currently available bacterial V_m_ fluorescent indicators can be categorized into two classes: (i) voltage-sensitive dyes and (ii) genetically encoded voltage indicators (GEVIs). Most members of the former class are Nernstian dyes, a group of cationic membrane-permeable dyes that distribute across the membrane according to the Nernstian equation (*16–17*). The typical Nernstian sensors for bacteria are TMRM (*18*) and ThT (*12*). However, the bacterial cell envelope is much more complex than the hypothetical free diffusion model. There are many factors that can influence the accumulation of Nernstian sensors in bacterial cells, such as the efflux activity or nonspecific adsorption on the membrane (*17*). Before use, careful calibration is required to obviate these factors (*17*). Additionally, in order to reach an equilibrium distribution of the dye inside and outside the cell, there must be a medium surrounding the bacteria with a constant concentration of the Nernstian dye, which prevents observation on a solid medium and may influence the physiology of bacteria during long-term tracking (*19*). Besides TMRM and ThT, carbon cyanine dyes such as DiOC_2_(3) also respond to bacterial V_m_. DiOC_2_(3) has been developed as a commercial bacterial V_m_ dye and can be used in a variety of bacteria (*20*). However, DiOC_2_(3) is cytotoxic and cannot be used to track the dynamics of V_m_ in living cells. In comparison with chemical dyes, GEVIs, which are endogenously expressed and not influenced by efflux pumps, offer a non-invasive and convenient method for V_m_ measurement. However, most GEVIs are designed for neurobiological experiments and their applications are limited to eukaryotic systems. So far, the only reported bacterial GEVI is PROPS (λ_em_ = 660-760 nm), which is a green-absorbing proteorhodopsin based mutant (*11*). Thus, it is highly desirable to develop an *in vivo* bacterial V_m_ sensor with high spatiotemporal resolution that can decipher the new aspects of bacterial electrophysiology.

Here, we have developed a new set of <u>**V**</u>oltage <u>**i**</u>ndicators for <u>**Bac**</u>teria, termed ViBac1 and ViBac2, by modifying the eukaryotic membrane voltage sensor ArcLight (*21*). We showed that these two sensors responded sensitively to both hyperpolarization and depolarization processes in bacteria, allowing single-cell recording of bacterial V_m_ dynamics *in vivo* with high temporal resolution down to a few hundreds milliseconds.

## Results

### Development and characterization of bacterial V_m_ sensor ViBac

Here we chose Arclight, a sensitive fluorescent protein voltage sensor for mammalian cells, for modification to develop a bacterial V_m_ sensors. As shown in Fig. 1A, ViBac1 was obtained by adding a short bacterial membrane targeting sequence (MTS) (*22*) to the N-terminus of ArcLight to facilitate localization at the inner membrane. ViBac2 was designed to enable crucial cross-cell V_m_ comparison by normalizing the influence of varying expression levels in different cells. A V_m_-inert protein (mCherry-L) was fused to the C-terminus of ViBac1 via a rigid linker, enabling the ratio of Arclight fluorescence intensity (I_g_) to that of mCherry-L (I_r_) to be utilized as an indicator of the relative V_m_ (Fig. 1A).

**Fig. 1.**
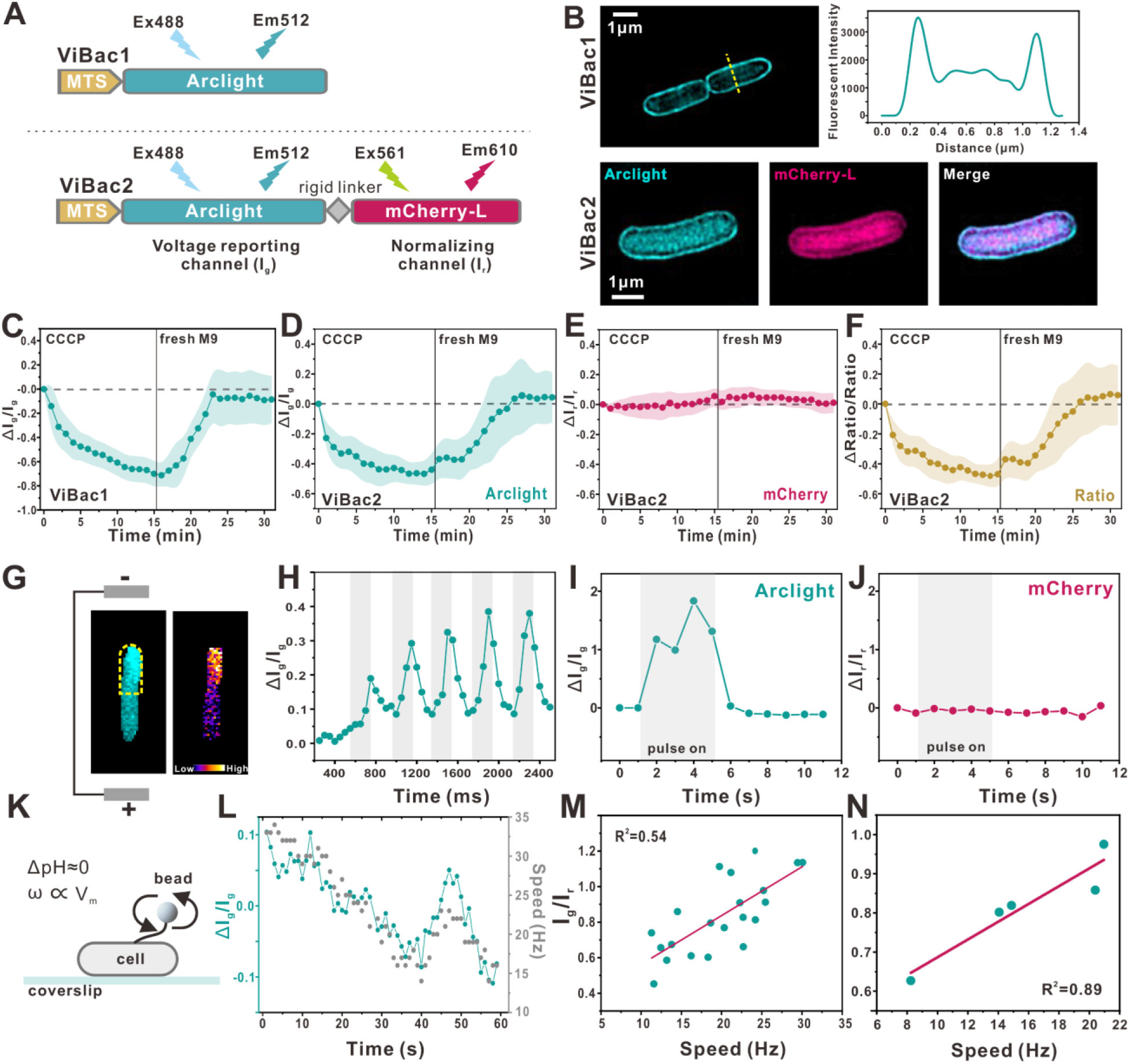
Development and characterization of the voltage sensors ViBac1 and Vibac2. (A) Schematics of ViBac1 and ViBac2 construction. ViBac1 is a green fluorescent protein (Ex=488 nm, Em=512 nm). ViBac2 is a double-channel fusion protein that emits green and red fluorescence. The fluorescence intensity in the green channel (I_g_, Ex=488 nm, Em=512 nm) responds to membrane voltage, and the red channel intensity (I_r_, Ex=561 nm, Em=610) is used to normalize protein expression. Thus, the fluorescence ratio (I_g_/I_r_) indicates the relative membrane voltage in *E. coli* cells. (B) Top left: SIM image of ViBac1 localization in *E. coli* cells. Top right: Fluorescent intensity along the short axis of a bacterial cell (the yellow line in the left panel). Bottom: SIM images of ViBac2 localization in *E. coli* cells (cyan: I_g_; red: I_r_). (C) Fluorescence response of ViBac1 to CCCP depolarization and subsequent M9 (0.4% glucose) recovery (N=15, error band: SD). (D-F) I_g_ (D), I_r_ (E), and fluorescence ratio (I_g_/I_r_) (F) of ViBac2 in response to CCCP depolarization and subsequent M9 recovery (N=15, error band: SD). (G) Spatially resolved change in fluorescence in a single cell subjected to ITV. (H) The dynamics of ViBac1 fluorescence in the half of the cell proximal to the negative pole of a microelectrode (circled region in A) during repeated short pulses of electrical stimulation (200 ms on, 200 ms off). (I-J) Changes in the I_g_ and I_r_ of cells expressing ViBac2 (half-cell proximal to the negative pole) that were subjected to ITV during a single long pulse of stimulation (4 secs). (K) Schematic diagram of beads assay. The external pH was carefully adjusted to ensure pH_i_ ≈ pH_o_. (L) Simultaneous measurement of ViBac1 fluorescence intensity and bead rotation speed. (M) The fluorescence ratio of ViBac2 plotted against flagellar rotation speed. Each point represents an individual cell. Red line: linear regression (number of cells=21). (N) Typical example of fluorescence ratio/flagellar rotation speed measurements from the same cell at different time points.

Structured illumination microscopy (SIM) revealed that both ViBac1 and ViBac2 anchor successfully onto the *E. coli* inner membrane (Fig. 1B) and has no significant impact on cell growth (Supplementary Fig. 1A-C). The emission spectra of ViBac1 and ViBac2 *in vivo* were identical to that of ArcLight (Supplementary Fig. 1D-E). We first validated the membrane voltage response of ViBac1 and ViBac2 using standard depolarization and hyperpolarization agents in population measurement experiments. In comparison with the existing bacterial voltage indicators (DiOC_2_(3), TMRM and PROPS), ViBac1 and ViBac2 consistently responded to V_m_ alterations with high sensitivity and had low toxicity (Supplementary Fig. 2, Supplementary Note 1). As shown in Fig. 1C, ViBac1 fluorescence decreased rapidly upon a 15-minute exposure to a depolarizing agent (CCCP). After the cells were replenished with fresh M9 medium (0.4% glucose), ViBac1 fluorescence increased and recovered to its initial level. The changes in ViBac2 I_g_ in response to CCCP and fresh M9 medium were similar to those of ViBac1, while the normalizing fluorescence (I_r_) was stable (Fig. 1D-F). The maximal variation of I_g_ exceeded 40% for both ViBac sensors, demonstrating a high sensitivity to voltage alterations under physiological conditions. To test whether the change in ViBac fluorescence was the result of fluctuation in internal pH, we expressed the pH-sensitive fluorescent protein pHmScarlet, which has an optimal pKa (7.4) and a high apparent Hill coefficient (1.1), in *E. coli* cells (*23*). As shown in Supplementary Fig. 3A-B, the intracellular pH of *E. coli* cells was stable during treatment with CCCP and fresh M9 medium.

To further confirm the voltage response of ViBac sensors, induced transmembrane voltage (ITV) was applied to cells expressing ViBac1 or ViBac2 through microelectrodes (Fig. 1G). Upon stimulation of the cells with repeated short pulses (200 ms on, 200 ms off), the ViBac1 fluorescence signal became oscillatory, and each increase in intensity corresponded to the onset of each pulse (Fig. 1H). For cells expressing ViBac2, when a single long pulse (5 secs) was applied, I_g_ rose rapidly and plateaued, while I_r_ was steady (Fig. 1I-J). In summary, ViBac sensors enabled single-cell recording of bacterial V_m_ dynamics in live cells with high temporal resolution down to just a few hundreds of milliseconds.

Next, we confirmed that the ViBac fluorescence ratio is proportional to V_m_ by using the bacterial flagellar motor (BFM) as a real-time nanoscopic “voltmeter” (*24, 25*). Flagellar rotation and the ViBac fluorescence ratio were observed simultaneously by attaching a polystyrene bead to specially modified “sticky” flagellar filament stubs of cells expressing ViBac1 or ViBac2 (Fig. 1K). We found that the fluorescence intensity of ViBac1 and bead (1.0 μm in diameter) rotation speed were strongly correlated with a Pearson correlation coefficient 0.81±0.16 (±SD, number of cases =11). Fig. 1L shows a typical case where the fluorescence intensity of ViBac1 and bead rotation speed fluctuated simultaneously (Pearson correlation coefficient =0.87). For ViBac2, as shown in Fig. 1M, a strong positive correlation (Pearson correlation coefficient=0.73, p=1.60×10^-4^) between the fluorescence ratio and bead (1.4 μm in diameter) rotation speed was observed (number of cells=21). Considering that the BFMs of different cells may be driven by varying numbers of molecular motor stator units, we focused our measurement on a single BFM (Fig. 1N). By long-term time-lapse recording, we observed a linear relationship between the ViBac2 ratio and bead rotation speed when V_m_ gradually dissipated upon starvation. Therefore, the ViBac2 fluorescence ratio faithfully reflects the V_m_ heterogeneity in a population of bacteria.

### Hyperpolarization spikes in *E. coli* V_m_ revealed by ViBac1

Next, we explored bacterial electrophysiology using our newly developed V_m_ sensors. The spontaneous depolarization spiking of exponential-phase *E. coli* expressing ViBac1 in M9 salts (imaging interval = 1 sec) was similar to that reported by Kralj *et al*. (*11*) (Supplementary Fig. 4A); however, in addition to transient depolarization, we observed hyperpolarization events (Fig. 2B). Surprisingly, when the M9 salts was replaced with PBS, the frequency and amplitude of the transient hyperpolarization events increased significantly (Fig. 2C-D), suggesting that most of the cells had entered a hyper-excited state. To quantitatively describe bacterial hyperpolarization events in different types of media, we defined a hyperpolarizing “spike” as a peak in fluorescence intensity that exceeded 20% of the baseline intensity, and “spiking cells” were defined as cells that showed more than 2 spikes over a time scale of 300 secs. As shown in Fig. 2E, in M9 salts, only 2.8% of *E. coli* cells (number of total cells =180) showed hyperpolarizing spikes. In contrast, in PBS, 73.4% of *E. coli* cells (number of total cells =169) showed hyperpolarizing spikes, with some cells exhibiting more than 25 spikes within 300 secs. The average spike amplitude of *E. coli* cells in PBS was 0.48 ± 0.27 (±SD, number of spikes = 1475), which was nearly twice that of cells in M9 medium (0.27 ± 0.06, number of spikes = 134). To better characterize the spiking events of *E. coli* cells in PBS, a shorter imaging interval (100 ms) was used. As shown in Fig. 2F, the typical shape of each hyperpolarizing spike resembled that of a neuronal action potential, with a rapid rise to the peak followed by a slower decay back to the baseline, yielding an average asymmetric factor of 2.99 ± 2.84 (±SD, number of spikes = 105). The full duration at half maximum of the hyperpolarizing spikes was 5.1 ± 3.5 secs (±SD, number of spikes = 105).

**Fig. 2.**
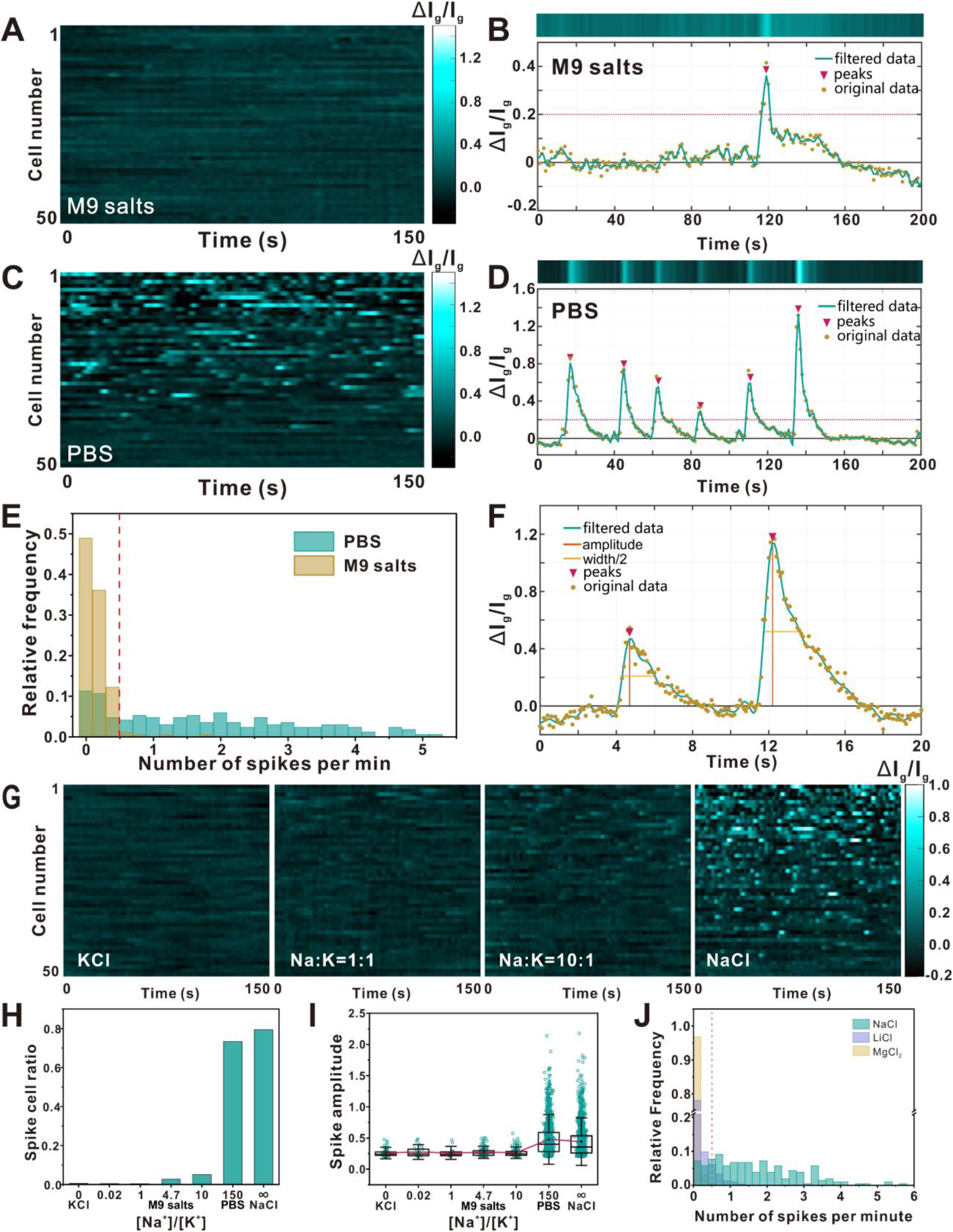
Distinct dynamics of *E. coli* membrane voltage in PBS and M9 salts. (A) Single-cell ViBac1 fluorescence time traces showing membrane voltage dynamics in M9 minimal medium recorded at 1-s intervals. (B) Typical transient hyperpolarization event of a cell in M9 salts. (C) Single-cell ViBac1 fluorescence time traces showing membrane voltage dynamics in PBS recorded at 1-s intervals. (D) Typical transient hyperpolarization events of a cell in PBS. (E) Histogram of the spike number per minute for every cell in M9 salts and PBS. (F) Typical spiking events of a cell in PBS recorded at 100-ms intervals. The asymmetric factor is defined as the distance from the center of the peak to the rearward-facing slope divided by the distance from the center to the forward-facing slope. All measurements were made at half-prominence. (G) Single-cell ViBac1 fluorescence time traces of cells in medium with a set Na^+^/K^+^ ratio. (H) The spiking cell ratio in medium with a set Na^+^/K^+^ ratio. (I) Distributions of spike amplitude in medium with a set Na^+^/K^+^ ratio. (J) Histogram of the spike number per minute for every cell in 300 mOsm/L NaCl, LiCl and MgCl_2_ solution.

### Ions trigger hyperpolarization spikes

To investigate the cause of the transient hyperpolarization and determine why cells kept in different types of media showed different hyperpolarization behavior, we compared the composition of M9 salts and PBS. Ammonium and anions (phosphate and chloride) were found to have little effect on spiking (Supplementary Fig. 4B-D, Supplementary Note 2), which led us to focus on the ratio of sodium and potassium ions (Na^+^/K^+^ ratio). As shown in Fig. 2G-I, the spiking frequency and amplitude increased significantly when the Na^+^/K^+^ ratio was increased under isosmotic conditions. In NaCl solution, 79.4% of cells were spiking cells (number of total cells = 209), while only one spiking cell was captured among the 177 cells that were recorded in KCl solution. We also verified that the spikes observed in NaCl did not arise from fluctuations in intracellular pH by applying the fluorescent pH indicator pHmScarlet (*23*) (Supplementary Fig. 3C, Supplementary Note 3).

To further examine the relative contributions of high [Na^+^] and low [K^+^] to the spiking behavior, we monitored the V_m_ dynamics of bacterial cells immersed in LiCl and MgCl2 solution. Very few spiking events were observed (Fig. 2J), indicating that Na^+^ is essentially required for the spiking behavior. We then tested the influence of [Na^+^] on the spiking behavior. As shown in Fig. 3A, the spiking behavior emerged gradually as [Na^+^] increased, and the ratio of spiking cells reached 73.82% when [Na^+^] reached 120 mM. When [Na^+^] was fixed at 120 mM, the spiking behavior was gradually suppressed as [K^+^] increased (Fig. 3B), and 30 mM [K^+^] was found to inhibit spiking. These results show that both high [Na^+^] and low [K^+^] are essential for hyperpolarization spikes in exponential-phase *E. coli* cells. Notably, the spiking behavior induced by a high Na^+^/K^+^ ratio was not unique to *E. coli*; we observed similar spiking behavior in *Salmonella typhimurium* and *Bacillus subtilis* that each expressed ViBac1 (Supplementary Fig. 5, Supplementary Table 1).

**Fig. 3.**
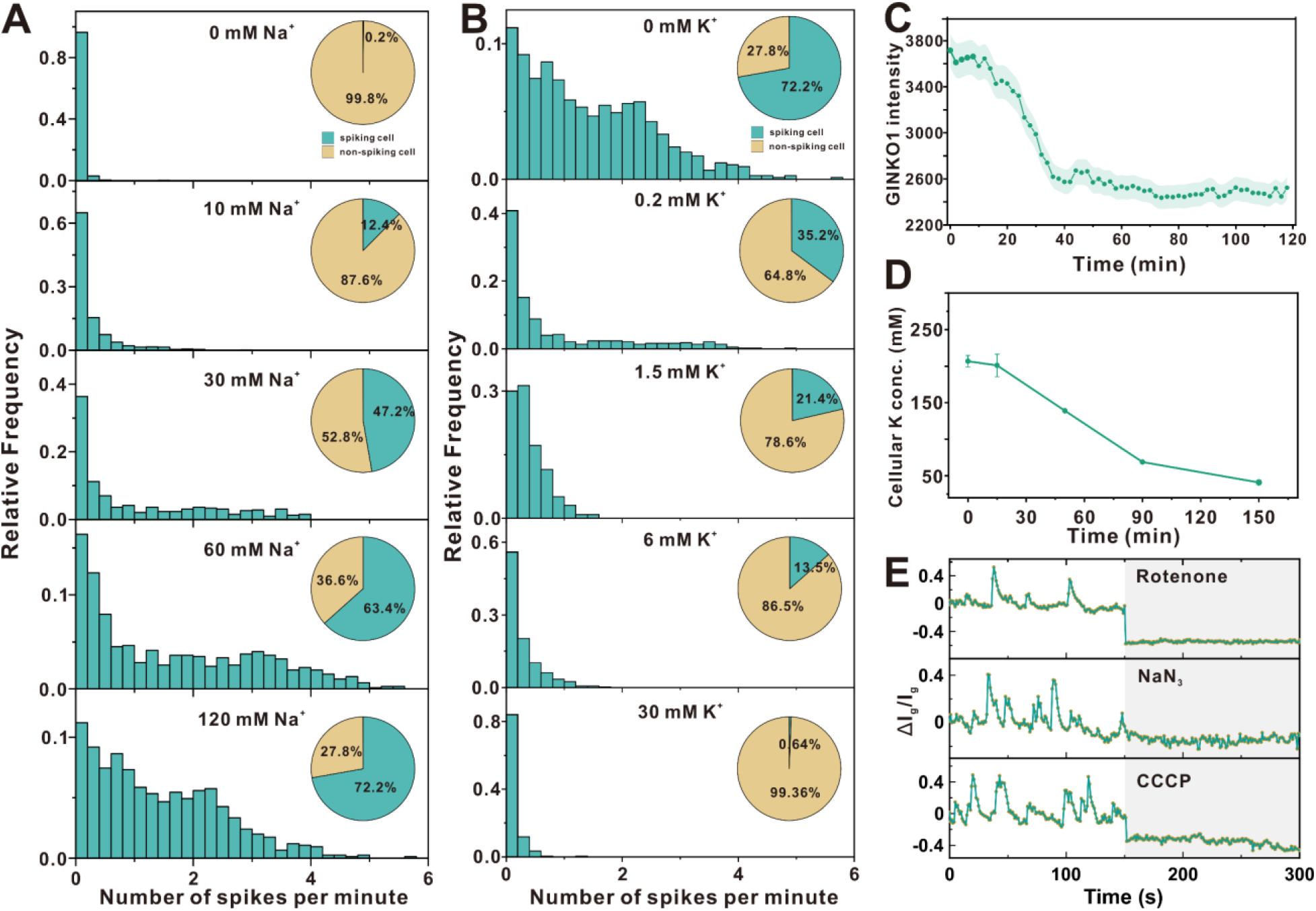
Excitability of exponential phase *E. coli* cells revealed by ViBac1. (A) Histogram of the spike number per minute for cells in solutions with different sodium concentrations. Sucrose was supplied to maintain an isotonic solution (300 mOsm/L) for each measurement. The inserted pie chart shows the ratio of spiking cells to non-spiking cells. (B) Histogram of the spike number per minute for every cell in solutions with 120 mM NaCl and different concentrations of potassium. Sucrose was supplied to maintain an isotonic solution (300 mOsm/L) for each measurement. (C) Fluorescence of GINKO1 in NaCl solution at a time scale of 2 hrs (N = 60, error band: SE). (D) Changes in cellular potassium concentration measured by ICP-OES after cells were transferred to NaCl solution. (E) The single-cell ViBac1 fluorescence before and after treatment with metabolic inhibitors (rotenone, NaN_3_ and CCCP).

### Hyperpolarization spikes are accompanied by K^+^ outflow

Changes in V_m_ usually involve the transmembrane movement of charged species. To investigate the electrochemical driver of each hyperpolarization spike, we focused on cytoplasmic K^+^, which is the most abundant cellular cation and plays a role in V_m_ regulation (*12*). By using the [K^+^] fluorescent indicator GINKO1 (*26*), we found that the cellular [K^+^] was quite stable at a time scale of 300 secs in NaCl solution (Supplementary Fig. 6). Due to the small membrane surface area and capacitance, the outflow of just a small fraction of cellular K^+^ ions would be expected to rapidly result in hyperpolarization (*27*). Thus, the cellular [K^+^] might not decline dramatically in a short period. By extending the observation time, we found that the cellular [K^+^] eventually decreased at a time scale of 2 hrs (Fig. 3C). The decrease in cellular [K^+^] was also quantitatively confirmed by ICP-EOS. As shown in Fig. 3D, the cellular [K^+^] concentration decreased in NaCl solution from 207 mM to 69 mM within 90 mins. Taken together, these results suggest that transient V_m_ hyperpolarization in bacterial cells immersed in NaCl solution is due to the outflow of cytoplasmic K^+^.

### Hyperpolarization spikes depend on the metabolic state

Since electrophysiology and metabolism are tightly coupled in bacteria, we examined how the spiking behavior responded to the metabolic state switch in *E. coli*. Inhibition of aerobic respiration using rotenone, NaN_3_ or CCCP all significantly suppressed the spiking of membrane voltage (Fig. 3E), indicating a tight link between aerobic respiration and V_m_ hyperpolarization. In addition, stationary-phase bacteria, in which aerobic respiration was down-regulated (*28*), did not show spiking behavior even in NaCl solution (Fig. 4A, left), and the cellular [K^+^] of stationary-phase cells decreased significantly less than that of exponential-phase cells after immersion in NaCl solution (Fig. 4A, right). To summarize, we have identified two distinct bacterial V_m_ states. When aerobic respiration is active (*e.g*. exponential-phase), V_m_ is excitable, especially with a high Na^+^/K^+^ ratio in the medium; in contrast, when aerobic respiration is down-regulated (*e.g*. stationary-phase), V_m_ shows no clear response to changes in ion concentrations.

**Fig. 4.**
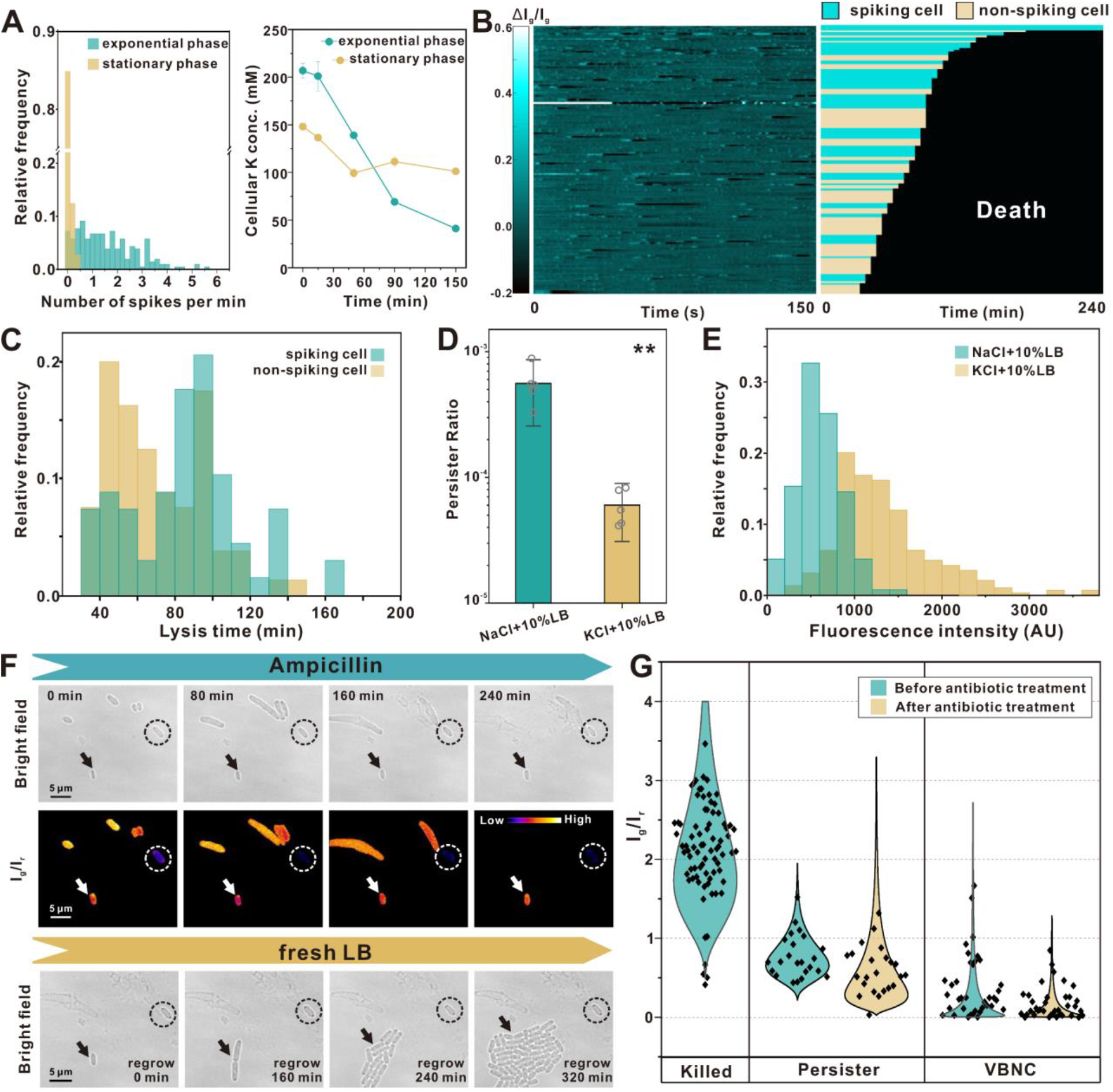
Role of membrane voltage dynamics and heterogeneity in antibiotic tolerance. (A) Left: Histogram of the spike number per minute for cells in the exponential and stationary phases. Right: Changes in the cellular potassium concentration of exponential-phase and stationary-phase cells measured by ICP-OES after transfer to NaCl solution. (B) Left: Single-cell ViBac1 fluorescence time traces in NaCl+10%LB acquired at the beginning of ampicillin treatment. Right: Single-cell traces showing the fate of each cell (corresponding to the left panel). Cyan stripes represent spiking cells. Brown stripes represent non-spiking cells. Black region marks the death of the tracked cells. (C) Distribution of the lysis time of the spiking and non-spiking groups. Representative images from a single replicate of two independent replicates are shown. (D) Persister ratio of cells in NaCl+10%LB and KCl+10%LB after 4 hrs of ampicillin treatment (unpaired Student’s t-test, error bar indicates SD, **p-value < 0.005). (E) Histogram of BOCILLIN accumulation in cells in NaCl+10%LB and KCl+10%LB. (F) Time-lapse images of stationary cells expressing ViBac2 during ampicillin killing and subsequent resuscitation (replacing the killing medium with fresh LB). Arrows indicate persister cells. Circle indicates a viable but non-culturable cell (VBNC). (G) Log-normal estimated distribution of the I_g_/I_r_ of ViBac2 in killed cells, persisters, and VBNCs before and after ampicillin treatment.

### Hyperpolarization spikes facilitate antibiotic tolerance

We next explored the relationship between V_m_ and bacterial fitness under antibiotic attack. First, we studied exponential-phase, in which V_m_ is in an excitable state. We compared the antibiotic tolerance of spiking cells and non-spiking cells by tracking their fates under treatment with 50 μg/mL ampicillin in NaCl with 10% LB for 240 mins. The supplementation with 10% LB did not significantly affect the spiking behavior (Supplementary Fig. 7). As shown in Fig. 4B-C, the lysis time of most spiking cells was significantly delayed in comparison with that of non-spiking cells (p=0.002 by Kolmogorov-Smirnov test). It is noteworthy that all persisters (cells that can regrow to form a colony after removal of antibiotics) were from the group of spiking cells, indicating that the electrical spiking of *E. coli* cells is associated with increased antibiotic tolerance. These findings were further confirmed using batch-culture persister assays. As shown in Fig. 4D, the persister ratio of cells in NaCl+10%LB was increased by an order of magnitude in comparison with that of cells in KCl+10% LB. Since V_m_ is an important component of PMF, we speculate that frequent V_m_ hyperpolarization might increase the bacterial efflux activity and therefore promote the drug tolerance. We measured the intracellular drug accumulation using a fluorescent β-lactam antibiotic BOCILLIN^™^ FL Penicillin (BOCILLIN). As shown in Fig. 4E, the average fluorescence intensity of BOCILLIN in cells immersed in NaCl +10%LB is about half of that immersed in KCl+10%LB, indicating that the antibiotics accumulated in spiking cells was substantially lower than that in non-spiking cells. Taken together, these results show that the V_m_ spiking state promotes bacterial survival under antibiotic exposure.

### ViBac2 revealed the heterogeneity of V_m_ and its link to antibiotic tolerance

Secondly, we studied stationary-phase cells, in which the V_m_ is in a resting state. Although the V_m_ was static, it displayed strong cell-to-cell heterogeneity, as evidenced by the wide distribution of ViBac2 ratios across each population of cells (C.V= 0.66, *N*=138). We then tracked the fate of cells with different V_m_ under 50 μg/mL ampicillin treatment for 240 mins. The antibiotic killing and subsequent resuscitation process of the bacterial population expressing ViBac2 were monitored in real-time at the single-cell level (Fig. 4F). According to the behavior of cells, we categorized them into three groups: ‘killed cells’ (lysed during antibiotic killing), ‘persisters’ (morphologically intact during antibiotic treatment and able to resuscitate after removal of antibiotics), and viable but non-culturable cells (VBNCs) (morphologically intact during antibiotic treatment but not able to resuscitate after removal of antibiotics). As shown in Fig. 4G, these three groups showed a diverse range of initial V_m_ before antibiotic treatment. Killed cells displayed the highest average V_m_, followed by persisters, and VBNCs possessed the lowest average V_m_. Of note, persisters were able to sustain their low V_m_ during antibiotic treatment, but, when fresh growth medium was replenished, the V_m_ of persisters was stable or slightly increased along with cell regrowth. Taken together, these results reveal the V_m_ heterogeneity in a bacterial population; drug-tolerant persisters showed an intermediate V_m_, whilst cells exhibiting high V_m_ were killed, and cells with ultra-low V_m_ were not resuscitative. However, we need to clarify that the higher V_m_ in some stationary phase cells, which still remains in the normal physiological range, is fundamentally different from the transient hyperpolarization observed in exponential-phase cells.

## Discussion

We reported the development and characterization of two GEVIs, ViBac1 and ViBac2, for monitoring bacterial V_m_ *in viv*o. ViBac1 is a single channel sensor whose fluorescence intensity is proportional to bacterial V_m_. ViBac2 is a two-channel sensor whose ratio of green fluorescence intensity to red fluorescence intensity is proportional to V_m_. ViBac2 enables comparison of V_m_ between different individual bacterial cells, reflecting population heterogeneity.

Using ViBac1, we first reported the frequent hyperpolarization spikes in bacteria. Our investigation further showed that these frequent hyperpolarization spikes in V_m_ are triggered by high [Na^+^] and low [K^+^] environments. The spikes are also observed in another Gram-negative bacteria *Salmonella typhimurium* and a Gram-positive bacteria *Bacillus subtilis* in high [Na^+^] and low [K^+^] environments using ViBac1, suggesting that this phenomenon is not unique to *E. coli*, but is likely to be a widespread phenomenon across different bacteria species. We found that the hyperpolarization spikes enhance the efflux activity and thereby promote bacterial antibiotic tolerance. It is noteworthy that simply adjusting the Na^+^/K^+^ ratio of the medium can lead to an order of magnitude difference in persister ratio, suggesting that ion concentration can strongly influence bacteria antibiotic tolerance. Na^+^ and K^+^ are common ions in liquid environments and it is possible that the distinct V_m_ patterns triggered by different Na^+^/K^+^ ratio are recognizing environments with different Na^+^/K^+^ ratio, such as the host’s intra-and extracellular media.

Meanwhile, we found that the frequent hyperpolarization spikes triggered by high [Na^+^] and low [K^+^] are closely related to the metabolic state of bacteria. When bacterial metabolism is down-regulated, the V_m_ becomes static and enters a resting state. Using ViBac2, we tested the antibiotic tolerance of a bacteria population with a highly heterogeneous V_m_ distribution. Previous studies have reported that a low V_m_ may promote antibiotic tolerance (*29–32*), but these results were obtained largely via bulk measurement. Our result revealed that cells with relatively low V_m_ prior to antibiotic treatment were more likely to be persisters, however, cells with ultra-low V_m_ were VBNCs. In our previous work, we have introduced the concept of “dormancy depth” to provide a unifying framework for understanding both the persisters and VBNCs (*33*). Persister cells are considered to be in shallow dormancy depth, while VBNCs are in a deeper dormancy. We speculate that V_m_ can serve as an indicator of the dormancy depth.

In summary, our findings demonstrate the potential of our newly developed voltage sensors for new explorations of bacterial electrophysiology. It is striking that bacterial cells exhibit electro-excitable and resting states, resembling the features of neurons. The mechanisms underlying fine regulation of bacterial V_m_ and how it relates to bacterial fitness and signal transduction remain to be explored, and future studies could evoke new strategies to combat bacterial drug tolerance.

## Methods

### Bacterial Strains

All strains and plasmids used in this study are indicated in Supplementary Table 2-3. Unless specified, the bacterial cells were cultured in Luria Broth (LB) at 37 °C, 200 rpm.

### Construction of ViBac1 and ViBac2

The original sequence of ArcLight (A227D) was a gift from Dr. Peng Zou. The construction of ViBac1 (*MTScrr-AcrLight*) was performed by adding a membrane targeting sequence (MTS) derived from *E.coli* EIIA^Glc^ protein (*22*) to the N-terminus of the original ArcLight. The *MTScrr-AcrLight* PCR product was inserted into the pBAD/myc-His A vector at the *Nco I* and *Hind III* sites by *in vitro* Gibson assembly. The pBAD::*ViBac1* plasmid was then transformed into BW25993, SYC12, SL1344 and 3610 cells, separately. The selected strains were designated JT04, SYC12-JT04, SL1344-JT04 and 3610-JT04, respectively.

To construct ViBac2 (*MTScrr-ArcLight-mCherry-L*), a modified form of the red fluorescent protein mCherry-L (*34*), in which the alternative translation start site was replaced (M9L), was connected to the C-terminus of Arclight with a rigid linker (LEAEAAAKALE). Similarly, the gene was cloned into the pBAD plasmid, then transformed into BW25993 and SYC12 cells. The selected strains were designated AM02 and SYC12-AM, respectively.

### Expression conditions of protein-based sensors

For ViBac1 and ViBac2 expression, an inducer (arabinose) was added to the culture medium at a final concentration of 0.002% (for exponential phase cells) or 0.05% (for stationary phase cells). After 2 hrs of induction, the arabinose was washed away. For experiments requiring the induction of ViBac1, cells were then ready for further observation. For induction of ViBac2, cells were incubated in “stabilizing medium” for 1.5 hrs after the removal of arabinose to ensure complete maturation. M9 salts were used as the stabilizing medium for exponential cells, and filtered supernatant from a parallel culture was used as the stabilizing medium for stationary cells.

For PROPS expression, cells were grown to the exponential phase (OD_600_≈0.3) in LB medium at 33 °C. An inducer (0.0005% arabinose) was added along with 5 μM all-trans retinal. Cells were harvested after 3 hrs of induction.

For pHmScarlet expression, BW25993 cells carrying the pBAD-pHmScarlet plasmid were grown to the early exponential phase, and induction was achieved by adding arabinose (0.001%) for 1 hr.

For GINKO1 expression, cells were grown to the early exponential phase, and induction was achieved by adding arabinose (0.005%) for 2 hrs.

### Effect of ViBac expression on cell growth

Overnight cell cultures were transferred at a ratio of 1:100 into fresh LB medium containing different concentrations of arabinose: 0% (control), 0.0001%, 0.0002%, 0.001%, or 0.01%. The growth curve of each group of cells (OD_600_) was monitored using a microplate spectrophotometer (SpectraMax M5, USA).

### Response to depolarization and hyperpolarization reagents

For experiments on ViBac1, ViBac2 and PROPS, exponential phase cells expressing certain voltage indicator proteins were prepared as described above and washed in PBS by centrifugation. Two aliquots were treated separately with 10 μM CCCP or 10 μM nigericin, and the third aliquot was used as the control. After 30 mins of incubation at 37 °C in the dark, cells were loaded on a gel-pad containing 2% low-melting-temperature agarose for microscopy.

For the application of voltage dyes (TMRM and DiOC_2_(3)), exponential-phase BW25993 cells (OD_600_≈0.3) were first resuspended in PBS. TMRM or DiOC_2_(3) was added to the medium at a final concentration of 100 nM or 30 μM respectively. Cells were then treated with 25 μM CCCP or 20 μM nigericin, and a third group was used as the control. After 30 mins of incubation at 37 °C in the dark, TMRM-stained cells were directly loaded onto a gel-pad for microscopy, while DiOC_2_(3)-stained cells were washed with PBS twice before they were loaded onto a gel-pad.

### Bright-field and fluorescence microscopy

Bright-field and epi-fluorescence imaging were performed on an inverted microscope (Zeiss Observer Z1, Germany). Illumination was provided by solid-state lasers (Coherent OBIS, USA) at various excitation wavelengths: 488 nm for ViBac1, ViBac2 (green channel), GINKO1, and DiOC_2_(3) (green channel); 561 nm for ViBac2 (red channel), TMRM, DiOC_2_(3) (red channel), and pHmScarlet; and 640 nm for PROPS. The fluorescence emission signal was recorded by an EMCCD camera (Photometrics Evolve 512, USA). Appropriate filter sets were selected for each fluorophore according to their excitation and emission spectra. For the observation of membrane potential dynamics using ViBac1 or ViBac2, the laser power output from the objective was set to 20-25 μW. An FCS2 flow cell chamber system with a temperature controller (Bioptechs) was applied for single-cell observation.

### Time-lapse recording of CCCP and subsequent M9 treatment

To monitor voltage dynamics with depolarizing or hyperpolarizing agents, cells expressing ViBac1 or ViBac2 were prepared as described above and suspended in M9 medium. Cells were attached to a coverslip pre-coated with poly-L-lysine and observed under a microscope at 37 °C. The initial V_m_ was recorded before the introduction of depolarizing agent CCCP. Next, PBS with 10 μM CCCP was injected into the flow cell chamber system for 15 minutes, while time-lapse recording of bright field and fluorescence images were performed (interval = 1 min, exposure time = 100 ms). Then, CCCP was washed away by supplementation with fresh M9 medium (0.4% glucose). The recovery period was recorded for an additional 15 minutes (interval = 1 min).

### Time-lapse recording of voltage dynamics with ViBac1 in different solutions

To monitor the voltage dynamics of cells in various salt solutions, cells expressing ViBac1 were suspended in the corresponding salt solution. The cells were then loaded onto a gel-pad containing 2% low melting temperature agarose. The gel-pad was prepared in the center of the FCS2 chamber as a small gel island and surrounded by flowing salt solution (volume of the liquid medium:volume of gel-pad = 20:1). The gel-pad was prepared with 0.5× salt solution to avoid hyperosmotic shock. The sample temperature was maintained at 37 °C by preheating the solution and tips to 37 °C. The sample was observed under a microscope at 37 °C. The exposure time was 100 ms and imaging interval was 1 s.

### Time-lapse recording of antibiotic killing and bacteria resuscitation

For the exponential phase, cells expressing ViBac1 were collected, washed two times with 150 mM NaCl solution and then loaded onto a gel-pad (90% 150 mM NaCl + 10% LB) containing 2% low melting temperature agarose. The cells were first imaged on the gel-pad surrounded by flowing medium to record the spiking events, after which they were exposed to 50 μg/mL ampicillin in 90% 150 mM NaCl + 10% LB medium for 4 hrs, and fresh growth medium LB was then flushed into the chamber to remove the antibiotic. The cells were monitored for another 16 hrs to observe resuscitation. Each sample was kept at 37 °C during the imaging process.

For the stationary phase, cells expressing ViBac2 were collected, washed two times with M9 medium, and then imaged on a gel-pad (90% M9 + 10% LB, 2% agarose) surrounded by flowing medium. The cells were first imaged in 90% M9 + 10% LB medium before antibiotic treatment. To record antibiotic-mediated killing and bacterial resuscitation, cells were exposed to 100 μg/mL ampicillin in 90% M9 + 10% LB medium for 4 hrs, after which fresh growth medium (LB) was flushed into the chamber to remove the antibiotic. The cells were monitored for another 16 hrs to observe resuscitation. Each sample was kept at 37 °C during the imaging process.

### Bead assay

Bead assays were performed on a custom-built inverted microscope equipped with an EMCCD camera (Andor iXon, UK) and a back-focal-plane interferometer. Illumination was provided by solid-state lasers (Coherent, OBIS, USA). The illumination wavelength was 488 nm for ViBac1 and ViBac2 (green channel), 561 nm for ViBac2 (red channel), and a 976 nm laser (Thorlabs, USA) was used for interferometer illumination.

Overnight-cultured SYC12-JT04 or SYC12-AM cells incorporating the sticky allele (*35*) were transferred into TB medium at 1:100 and cultured for 1 hr at 30 °C, 200 rpm. Arabinose was introduced to the medium at a final concentration of 0.002%, and the cells were incubated for another 2 hours at 30 °C. Later, cells were washed twice with motility buffer (MB) and resuspended in MB for an extra 40 minutes of incubation. Flagella were then sheared by repeatedly passing the cell suspension through the needle of a 1-mL syringe (*36*). Sheared cells were attached to a custom-made flow channel pre-treated with poly-L-lysine. Subsequently, carboxyl latex beads (Invitrogen) were added to the flow channel and incubated for 10 min. Unattached beads were then removed by allowing MB to flow gently through the channel.

The slides were mounted onto the microscope after sample preparation. The rotation of flagellum-attached beads was recorded at 10 kHz using the back-focal-plane interferometer. Simultaneously, the fluorescence signals were recorded using epi-fluorescence imaging. The flagellar rotor speed was calculated by processing the bright-field image sequences using a custom Python program.

### Structured Illumination Microscopy (SIM)

SIM images were acquired on a DeltaVision OMX SR imaging system (GE Healthcare, USA) equipped with a 100× oil-immersion objective (NA 1.49) and EMCCD, which achieved imaging of samples at approximately 120 nm lateral resolution. Laser lines at wavelengths 488 and 561 nm were used for excitation. The microscope was routinely calibrated with 200 nm diameter fluorescent microspheres. Images were reconstructed with the softWoRx 5.0 software package.

### Fluorescence spectral imaging

To measure the fluorescence spectra of ViBac1 and ViBac2 in living cells, cells expressing ViBac1 or ViBac2 were placed on an M9 gel-pad and observed using a laser scanning confocal microscope (Nikon A1R si+, Japan) equipped with a 100×/1.4 NA oil immersion objective lens. Fluorescence images were obtained using a laser scanning confocal microscope (Nikon A1R si+, Nikon Instruments, Japan) equipped with an APO 40×1.25 NA water immersion objective. The fluorescent spectra of ViBac1 (under 488 nm excitation) and ViBac2 (under 488nm and 561 nm excitation) were obtained using a 32-PMT spectral detector and then used for spectral unmixing. Spectral images were unmixed using NIS-elements AR software (Nikon, Japan) to obtain images of ViBac1 and ViBac2. All images were captured and analyzed using NIS Elements AR software (Nikon, Japan).

### Microelectrode assay

The microelectrode slides were generous offerings from Dr. Shuichi Nakamura. The distance between the electrodes on the surface of the bottom glass slide is around 60 μm. Vacuum grease was first applied to a glass slide using a syringe to form a rectangle. In order to thin the grease, the grease was pressed under another slide with finger pressure. After two rounds of such transfer, the thin grease layer was finally transferred to the electrode side of the microelectrode slide. Cells expressing ViBac1 or ViBac2 were attached to a poly-L-lysine-coated coverslip and washed with a sufficient amount of deionized water to completely remove the remaining ions. Then the coverslip was placed on top of the microelectrode slide with thin grease (facing the electrode side). The depth of the channels was estimated to be around 10 μm. Silver wires were used to connect the microelectrode slide to the voltage-generating circuit. The voltage pulses (3.75V) generated by a voltage stimulator (Grass Technologies, S88) and an isolator (Grass Technologies, SIU5) were applied to the attached cells. The voltage pulses were accompanied by continuous recording of the cellular fluorescence, and cells with long axes parallel to the electric field were selected for analysis.

### Antibiotic treatment and persister cell counting

Bacterial cultures (BW25993) were first grown to the mid-exponential phase. For antibiotic killing in salt solutions with 10% LB, log-phase cultures were first washed with corresponding salt solutions and diluted at 1:2 in salt solutions containing 10% LB and a lethal concentration of antibiotics (100 μg/mL ampicillin). Cells were then incubated in a shaker (200 rpm) for 4 hours at 37 °C. After killing, samples were then removed, diluted serially in PBS buffer and spotted on LB agar plates for overnight culturing at 37 °C. Colony formation unit (CFU) counting was performed the next day. CFU experiments were performed with five replicates.

### BOCILLIN accumulation measurement

For intracellular fluorescent antibiotic accumulation measurement, two aliquots of the LB-cultured bacterial cells (BW25993) were washed with 150 mM NaCl+10%LB or 150 mM KCL+10%LB twice by centrifugation and resuspended in the respective solutions. Fluorescent antibiotic BOCILLIN was added at a finial concentration of 20 μg/mL. Then the cells were incubated at 37°C in dark for 30 min. After that, the BOCILLIN was removed by centrifugation and cells were resuspended in the respective solutions. The cells were used directly for epifluorescence microscopy.

### Inductively Coupled Plasma Optical Emission Spectroscopy (ICP-OES)

The bacterial cells (BW25993) were collected by centrifugation (100 mL bacterial culture, 4000 rpm, 10 mins) and the samples were then subjected to a microwave digestion system (Milestone UltraWAVE ECR, Italy) for digestion. ICP-OSE (Leeman Prodigy 7, USA) was used to determine the potassium content. The intracellular potassium concentration was determined by normalizing the potassium levels by cell number per wet weight and single cell volume.

### Data analysis

#### (1) Analysis of spiking cells

In order to quantitatively analyze the spiking behavior of bacterial cells in various conditions, a semi-automatic image processing workflow was applied to the raw image sequences based on MATLAB (R2019a) and the ImageJ plugin MIJI. Firstly, the illumination field of the fluorescence channel was adjusted for each frame based on the image of a uniformly distributed green fluorescence protein solution. Then registration was applied to the time-lapse image sequences to correct minor displacements (*37*). Next, *Trainable WEKA Segmentation* (*38*) machine learning algorithm was applied to the images to create segmented masks for each cell, and the mean fluorescence intensity of every single cell at each frame was collected for further analysis.

To analyze and quantify the spiking behavior, the fluorescence time traces of each cell were first normalized based on the median fluorescence:

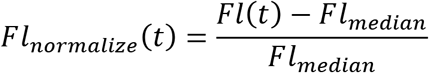

The normalized fluorescence time traces were processed through a low-pass filter to remove high frequency noise. Subsequently, spiking events were identified based on a peak with prominence greater than 0.2 (maximum instrumental error around 10%). The amplitude and duration of the spikes were also subtracted for further analysis.

#### (2) Analysis of stationary cells

To analyze the fluorescence ratio of ViBac2-expressing stationary phase cells, the background fluorescence was first subtracted from each image. Threshold-based segmentation was then performed based on the red fluorescence channel to recognize the outline of each bacteria cell. The fluorescence ratio (I_g_/I_r_) of every segmented cell was calculated by dividing the mean fluorescence intensity of the green (488 nm) and red (561 nm) channels. The pseudo-color images were generated by making a pixel-by-pixel fluorescence division of the background-subtracted green and red channel image within the regions that were recognized as cells, while the pixel value in the regions without cells was kept as zero.

## Data availability

Any data that support the findings of this study beyond what is included in the Supplementary Information are available from the corresponding author upon request.

## Acknowledgments

This work was financially supported by the National Science Fund for Distinguished Young Scholars (T2125002), Beijing Natural Science Foundation (JQ18019) to F.B., by BBSRC (BB/W000555/1) to M.C.L., by the Royal Society and NFSC (IEC\NSFC\191406) to F.B. and M.C.L, and by China Postdoctoral Science Foundation (2019M660307) to X.J., by the Ministry of Science and Technology of the Republic of China under contract (No. MOST-109-2628-M-008-001-MY4) to C.L.. We thank Dr. Yulong Li for technical assistance in microelectrode assay and the National Center for Protein Sciences at Peking University in Beijing, China for assistance with SIM imaging.

## Author contributions

F.B., X.J., and X.Z. conceived the study; X.J., X.Z., X.D., T.T., C.T., X.L., and X.C. performed the experiments; X.J. and X.Z. performed the data analysis; F.B., X.J., X.Z., C.L., and M.C.L wrote the manuscript.

## Additional information

**Supplementary information** is available for this paper.

**Materials & Correspondence** should be addressed to F.B.

