## Supplementary Materials for "Sensitive bacterial V_m_ sensors revealed the excitability of bacterial V_m_ and its role in antibiotic tolerance"

**This PDF file includes:**

Supplementary Figs. 1 to 7
Supplementary Tables 1 to 3
Supplementary Note 1 to 3

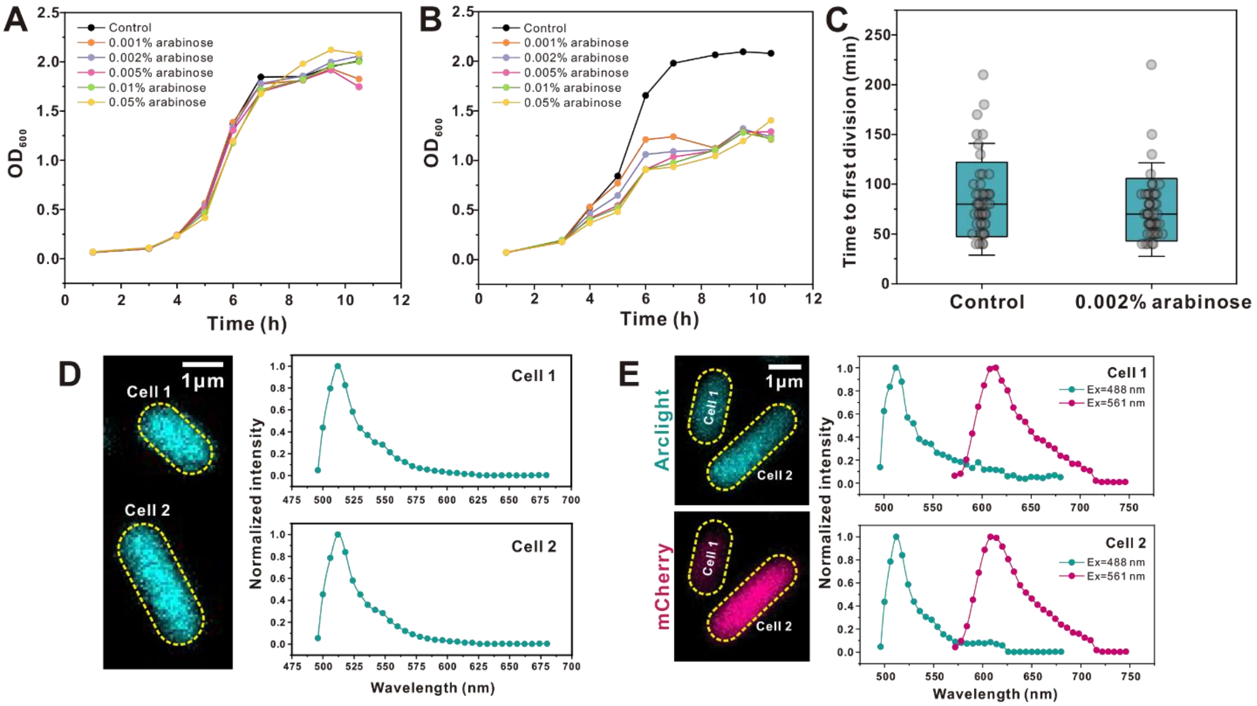


**Supplementary Fig. 1 Growth and emission spectrum of cells expressing ViBac1 or ViBac2**

(A) Growth curves of cells expressing ViBac1 with different arabinose concentrations. (B) Growth curves of cells expressing ViBac2 with different arabinose concentrations. (C) Time to the first division of cells with and without transient induced (0.002% arabinose, 2 hrs) ViBac2 expression under a microscope. (D) Spectral imaging reveals the emission spectrum of ViBac1 in living *E. coli* cells. Excitation=488 nm, wavelength resolution=6 nm. (E) Spectral imaging reveals the emission spectrum of ViBac2 in living *E. coli* cells. Excitation=488 and 561 nm, wavelength resolution=6 nm.

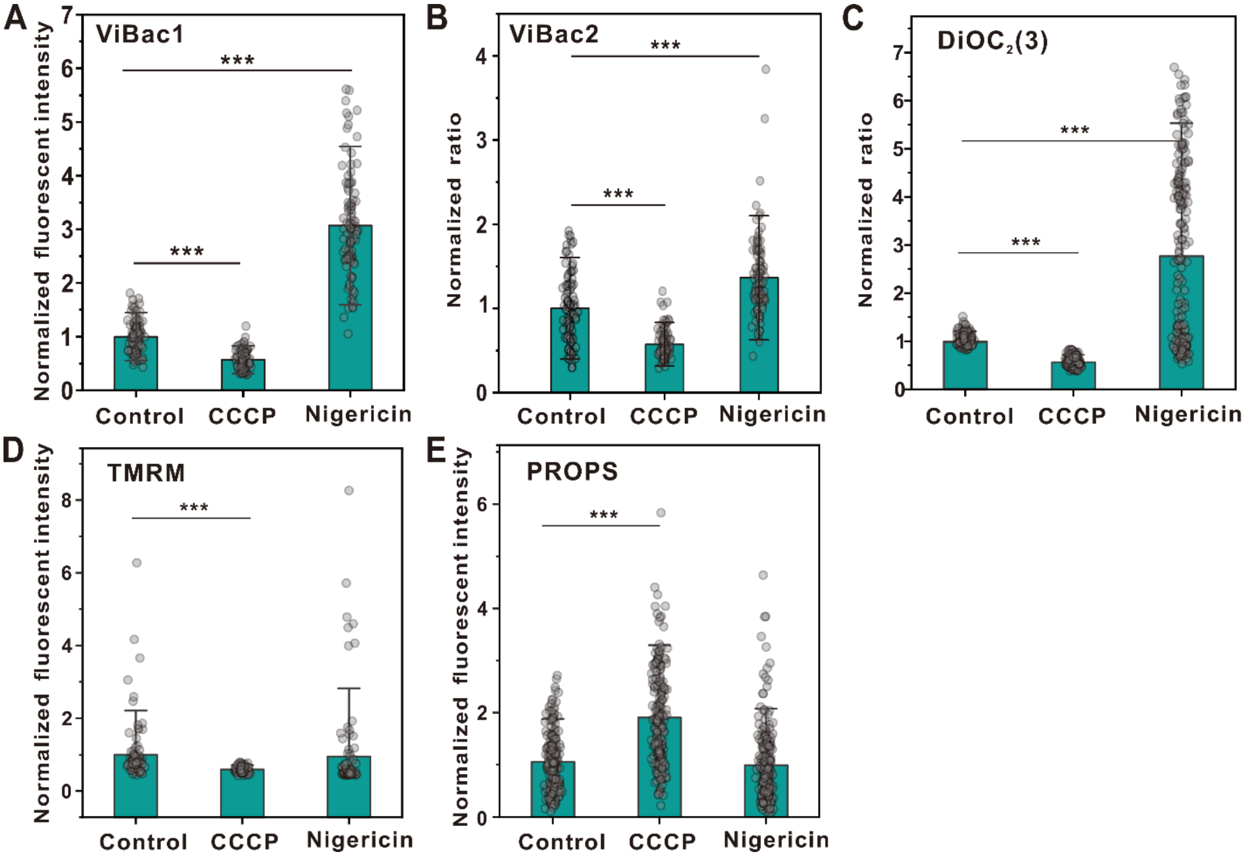


**Supplementary Fig. 2 Comparison of ViBac1 and ViBac2 with other established bacterial membrane voltage sensors.**

(A) ViBac1; (B) ViBaC2; (C) DiOC2(3); (D) TMRM; (E) PROPS

(unpaired Student's t-test against control, error bar indicates SD, *p value < 0.05; **p value < 0.005; ***p value < 0.0005)

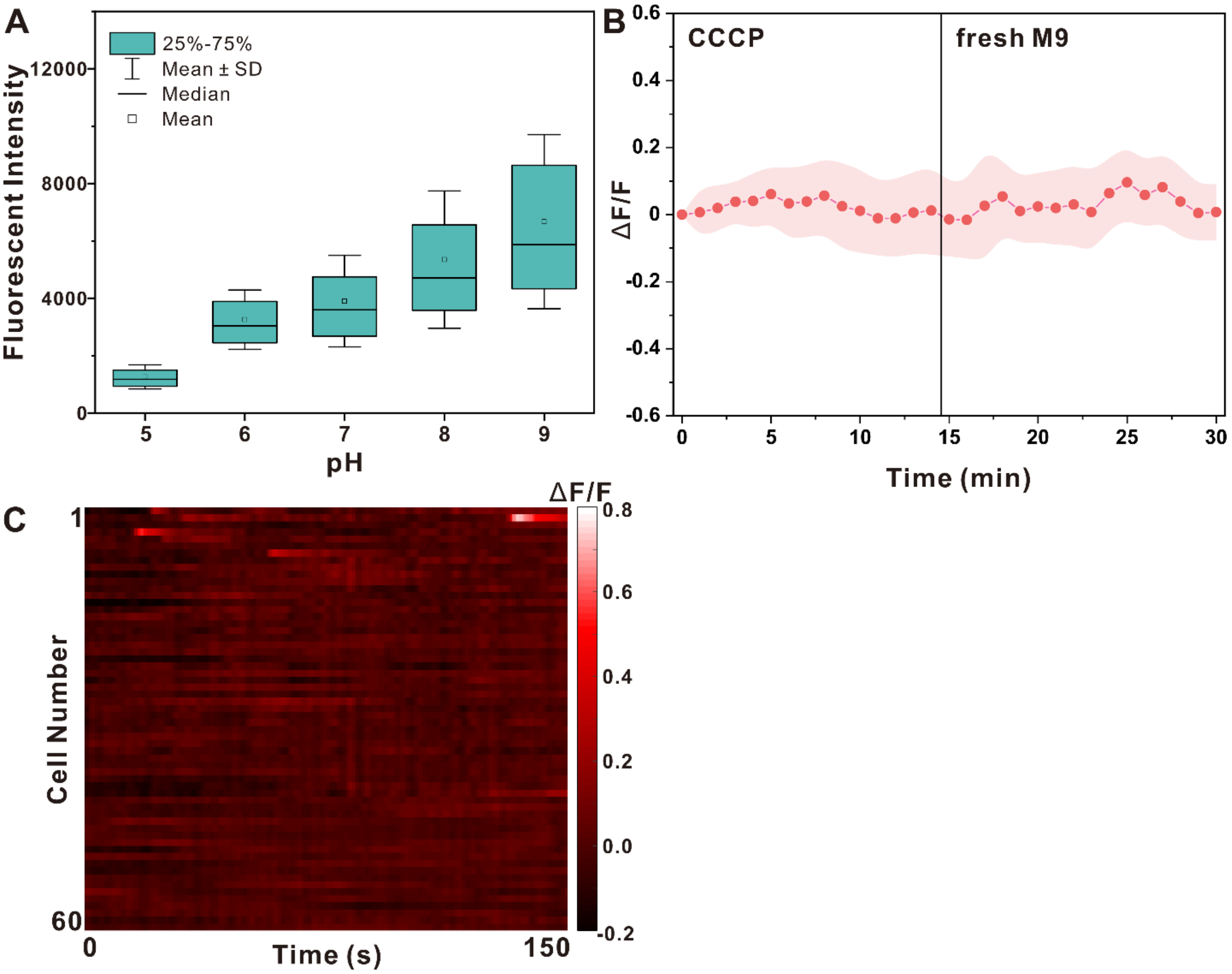


**Supplementary Fig. 3 The cellular pH is stable in NaCl solution.**

(A) Fluorescent intensity of cells expressing pHmScarlet after incubation in medium with varying pH containing protonophore. (N=1000); (B) Fluorescence response of pHmScarlet to CCCP depolarization and subsequent M9 (0.4% glucose) recovery treatment (N=15, error band: SD). (C) Single-cell pHmScarlet fluorescent intensity time traces of cells in 150 mM NaCl solution.

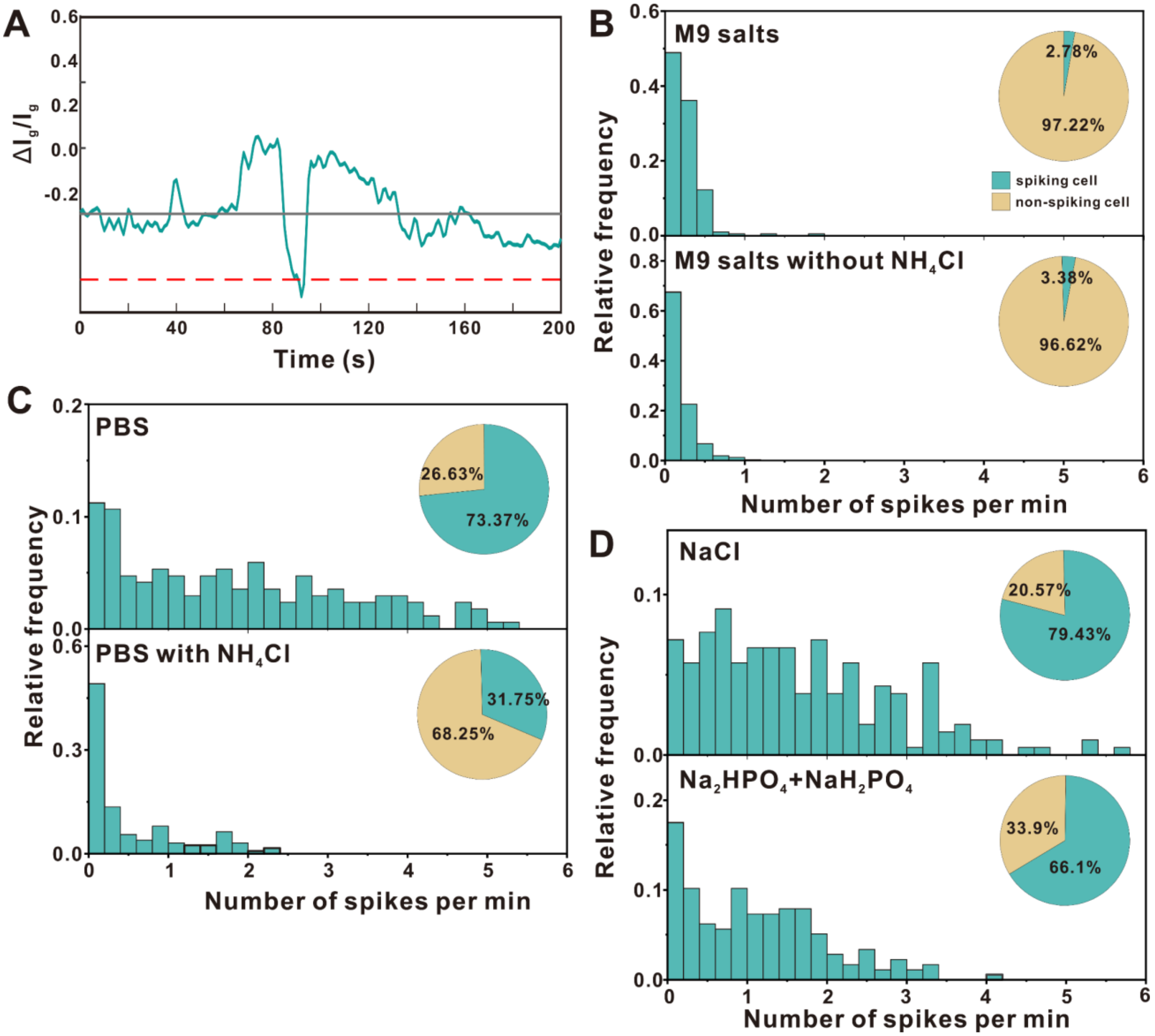


**Supplementary Fig. 4 Membrane voltage dynamics of *E. coli* cells in different solutions**

(A) Typical transient depolarization event of cells in M9 salts. (B-D) Histogram of the number of spikes per minute in different solutions. The inserted pie chart shows the ratio of spiking cells to non-spiking cells.

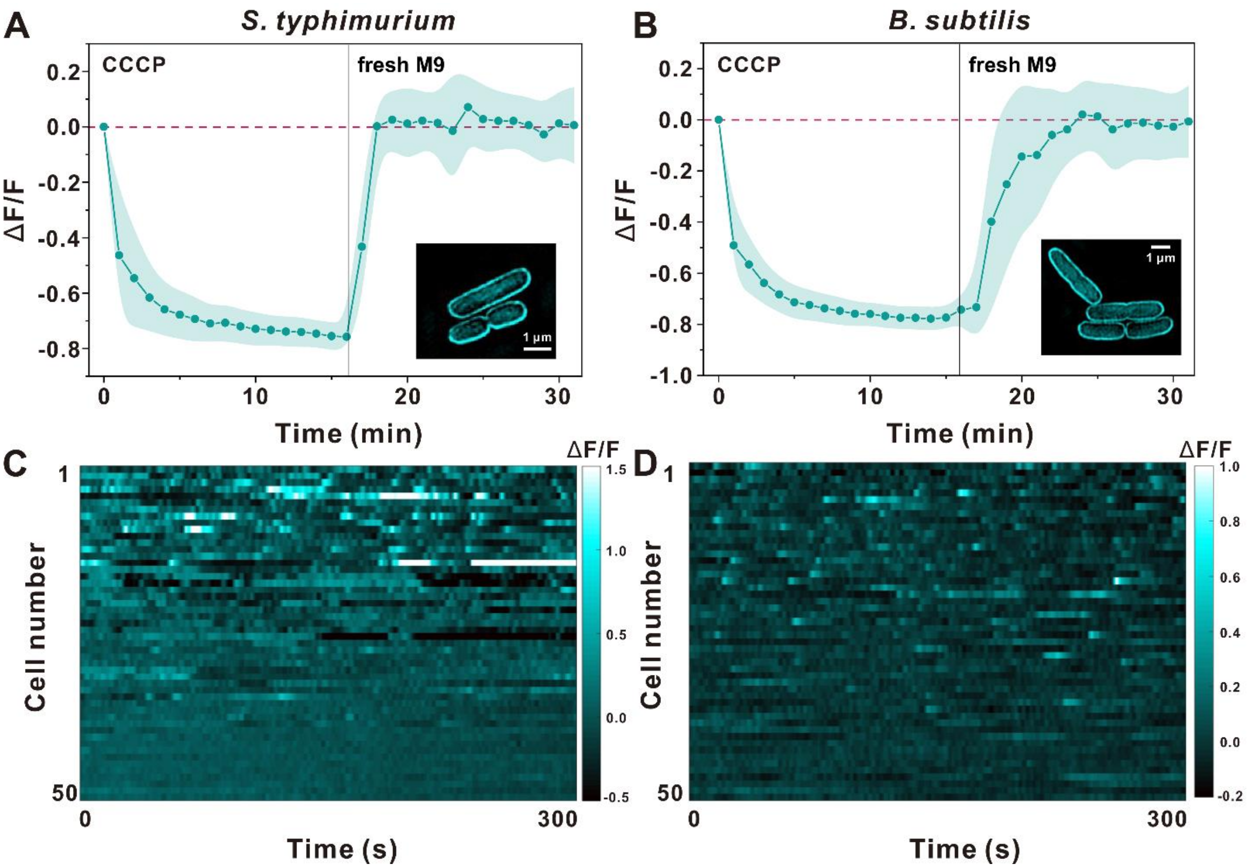


**Supplementary Fig. 5 Spiking behavior of *S. typhimurium* and *B. subtilis***

(A-B) Fluorescence response of ViBac1 in *S. typhimurium* (A) and *B. subtilis* (B) to CCCP depolarization and subsequent M9 (0.4% glucose) recovery treatment (N=15, error band: SD). Insert: SIM image of ViBac1 localization in *S. typhimurium* and *B. subtilis*. (C-D) Single-cell ViBac1 fluorescence time traces showing the membrane voltage dynamics of *S. typhimurium* (C) and *B. subtilis* (D) in saline solution recorded at 1-s intervals.

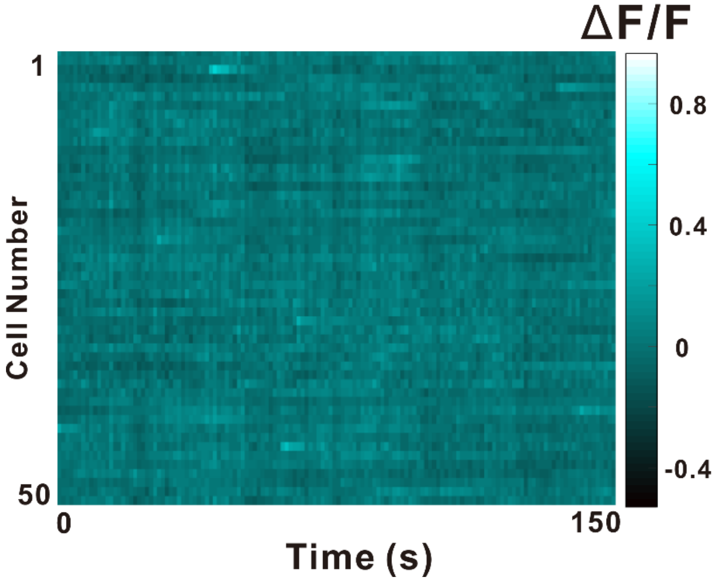


**Supplementary Fig. 6 Single-cell GINKO1 fluorescence time traces showing membrane voltage dynamics in 150 mM NaCl solution recorded at 1-s intervals.**

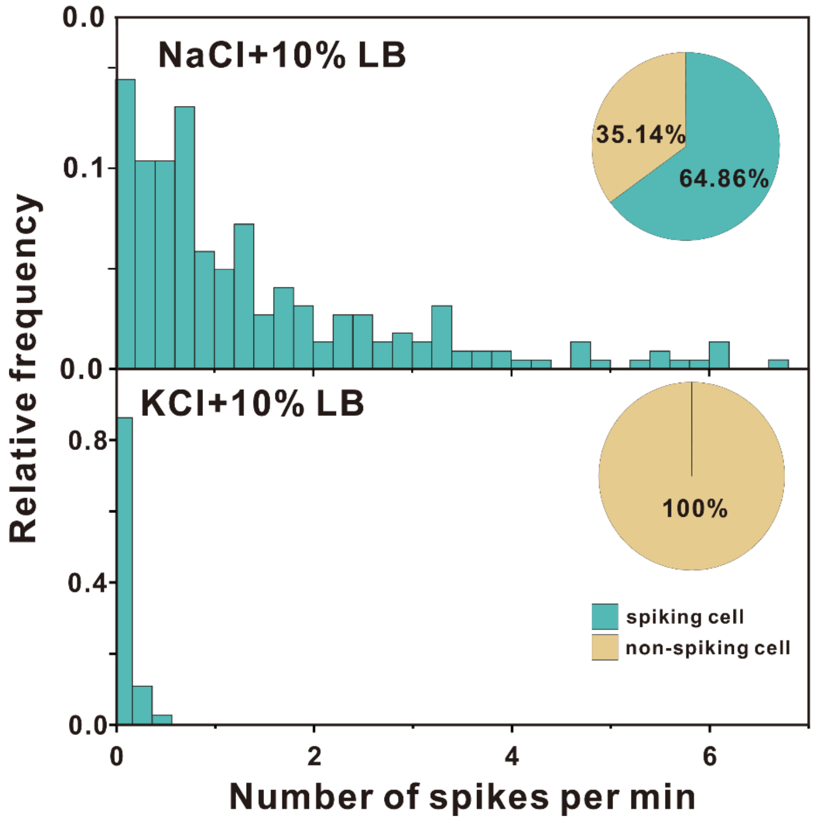


**Supplementary Fig. 7 Histogram of the number of spikes per minute of cells immersed in NaCl + 10%LB (top) or KCl + 10%LB (bottom). The inserted pie chart shows the ratio of spiking cells to non-spiking cells.**

**Supplementary Table 1. Spiking parameters of different strains**

| | ***E. coli*** | ***S. typhimurium*** | ***B. Subtilis*** |
|---|---|---|---|
| Spike cell ratio | 74% | 73% | 52% |
| Spike amplitude | $0.54 \pm 0.03$ | $0.45 \pm 0.01^{***}$ | $0.33 \pm 0.01^{***}$ |
| Spike duration (sec) | $3.36 \pm 0.23$ | $4.84 \pm 0.14^{***}$ | $5.82 \pm 0.31^{***}$ |
| Asymmetric factor | $2.24 \pm 0.11$ | $2.32 \pm 0.08$ | $2.98 \pm 0.15^{*}$ |

(Mann-Whitney test: *p<0.05,**p<0.01,***p<0.001)

**Supplementary Table 2. Strains used in this study**

| Strain | Background | Genotype |
|---|---|---|
| BW25993 | *Escherichia coli K12* | *Δ(araD-araB)567, $\lambda^-$, rph-1, Δ(rhaD-rhaB)568, hsdR514* |
| SYC12 | *Escherichia coli* RP437 | ΔfliC::fliC$^{st}$ |
| SL1344 | *Salmonella enterica subsp. enterica serovar Typhimurium str. LT2* | wild type |
| NCIB 3610 | *Bacillus subtilis subsp. Subtilis str. 168* | wild type |

**Supplementary Table 3. Plasmids used in this study**

| Plasmid | Gene | Induction condition |
|---|---|---|
| pBAD-ViBac1 | pBAD::MTScrr*-ArcLight | Arabinose 0.002% (log phase)<br>Arabinose 0.05% (stationary phase) |
| pBAD-ViBac2 | pBAD::MTScrr*-ArcLight-rigid linker†-mCherry(A9L) | Arabinose 0.05% (stationary phase) |
| pBAD-PROPS | pBAD::PROPS | Arabinose 0.0005% |
| pBAD-pHmScarlet | pBAD::pHmScarlet | Arabinose 0.001% |
| pBAD-GINKO1 | pBAD::GINKO1 | Arabinose 0.005% |

* MTScrr DNA sequence:
ATGGGTTTGTTCGATAAACTGAAATCTCTGGTTTCCGACGACAAGAAGGATACC
† Linker DNA sequence: AAGGCTGCACTAGAAGCCGAGGCAGCAGCAAAGGCCCTAGAA

### Supplementary Note 1

#### Comparison with other established bacterial voltage indicators

To confirm that our ViBac sensors could faithfully report $V_m$ and to address potential applications, we compared the voltage responses of the ViBac sensors and other established bacterial $V_m$ sensors. We chose the Nernstian dye TMRM, the membrane dye $DiOC_2(3)$, and the bacterial GEVI PROPS for comparison. As shown in Extended Data Fig. 2, ViBac sensors responded to $V_m$ alterations in the same manner as the commercial bacterial $V_m$ dye $DiOC_2(3)$. TMRM was sensitive to depolarization reagent CCCP, but not to hyperpolarization reagent nigericin. For TMRM staining, a fraction of the cells always had very weak fluorescence intensity, which was only 50% higher than the background, which might have been caused by poor loading of TMRM. The bacterial GEVI PROPS becomes significantly brighter upon depolarizing treatment. However, PROPS did not respond to hyperpolarization reagents, primarily due to its low baseline fluorescence. Due to its low fluorescence, PROPS requires high illumination power to ensure excitation, and further increasing the intensity of excitation could potentially damage the cells. Therefore, PROPS is not applicable for the observation of bacterial $V_m$ hyperpolarization.

The results above demonstrated that our ViBac sensors could faithfully respond to membrane voltage alterations with high sensitivity. The high temporal resolution (Fig. 1G-J) and low toxicity (Extended Data Fig. 1A-B) of our ViBac sensors are significant advantages in applications that require monitoring of the dynamics of bacterial $V_m$ *in vivo* at a single-cell level.

### Supplementary Note 2

#### Comparison of the formula of PBS and M9 salts

To investigate the cause of the different dynamics of $V_m$ in PBS and M9 salts, we compared the formula of PBS and M9 salts. As shown in the table below, the composition of the two solutions differ in three points: the $Na^+/K^+$ ratio, the presence of $NH_4^+$, and the anionic composition. As shown in Fig. 2, adjusting the $Na^+/K^+$ ratio changed the spike status significantly. We also investigated the influence of $NH_4^+$ and anionic on spike status.

**Comparison of the ion composition of PBS and M9**

| ion | Concentration in PBS (mM) | Concentration in M9 salts (mM) |
|---|---|---|
| $K^+$ | 1.06 | 22 |
| $Na^+$ | 161.11 | 103.9 |
| $NH_4^+$ | -- | 18.7 |
| $H_2PO_4^-$ | 1.06 | 22 |
| $HPO_4^{2-}$ | 2.97 | 47.7 |
| $Cl^-$ | 155.17 | 27.2 |

After removing the $NH_4^+$ from M9 salts, we found that there was no increase in spiking cell ratio (Extended Data Fig. 4B). After adding $NH_4^+$ to PBS, the spike frequency was reduced (Extended Data Fig. 4C). However, the spiking cell ratio was still around 10-fold higher than that observed in M9 salts. As shown in Extended Data Fig. 4D, the phosphate ions did not affect the spike status of $V_m$ in the $Na^+$-dominant solution. In addition, supplementing nutrients in sodium chloride solution had no significant effect on the spiking behavior (Extended Data Fig. 7), suggesting that the spiking behavior was not due to starvation.

**Supplementary Note 3**

**Cellular pH measurement using pHmScarlet**

To determine whether the spikes arose from cellular pH fluctuation, a pH-sensitive red fluorescent protein, pHmScarlet (*23*), was expressed in *E. coli*. To verify that pHmScarlet in *E. coli* can respond to the cellular pH, we incubated cells expressing pHmScarlet in medium containing protonophore (40 mM potassium benzoate and 40 mM methylamine hydrochloride), allowing us to modulate cytoplasmic pH through adjustment of the pH of the medium. After 30 mins of incubation at 37 °C, the cells were imaged under a fluorescence microscope. As shown in Extended Data Fig. 3A, the fluorescent intensity of pHmScarlet increased as the pH of the medium increased. We then observed dynamics of the pHmScarlet signal in NaCl solution. As shown in Extended Data Fig. 3C, the fluorescence fluctuated slightly, with a mean amplitude ($\Delta F/F$) of 0.28 ± 0.11. By assuming that the baseline fluorescence intensity represents a normal cellular pH (7.5), this amplitude was determined to correspond to a change in ($\Delta$pH) of 0.27 ± 0.10. Although there was a small group of cells showed a greater change in pH, the duration of the fluctuation in the pHmScarlet signal was significantly different from that of the spike in the ViBac1 signal. The mean fluctuation duration was 26.22 ± 11.77 secs for pHmScarlet and 5.1 ± 3.5 secs for ViBac1. These results demonstrate that the spikes in ViBac1 fluorescence did not arise from cellular pH fluctuation.
